# Exploratory Assessment of Preconception Phthalate Exposure on Fertility and Offspring Health in Mice

**DOI:** 10.1101/2025.08.25.671855

**Authors:** Maryam Afghah, Ansley C. Elkins, Paige C. Powell, Mary E. Mulligan, Mary C. Boland, Alexandra P. Suggs, Melissa A. Walker, Zachary J. Padgett, Kylie D. Rock

## Abstract

Infertility affects 10–15% of couples worldwide and is increasingly attributed to environmental exposures, particularly endocrine-disrupting chemicals (EDCs) such as phthalates. While gestational exposures are well studied, little is known about how preconception exposures influence fertility and offspring health. Phthalate exposure altered estrous cyclicity, with dams spending more time in proestrus and less in metestrus but did not significantly impact implantation or litter size. At E14.5, exposed fetuses exhibited increased bodyweights, accompanied by an expansion of the placental junctional zone in males. Altered bodyweight persisted into adulthood, however adulthood offspring displayed a reduction in bodyweight. RNA-sequencing revealed widespread transcriptional reprogramming in female placentas (518 DEGs) affecting immune regulation, steroid metabolism, and extracellular matrix remodeling, while male placentas exhibited only 9 DEGs but showed structural alterations. In offspring livers, transcriptomic shifts were sex-specific: females displayed downregulation of metabolic and immune genes (e.g., *Cyp7a1, Mthfr, H2-DMB1*), while males showed upregulation of immune and signaling genes (e.g., *Elf4, Adm2, Ly6a*). Collectively, these findings demonstrate that preconception phthalate exposure induces subtle but biologically meaningful maternal endocrine disruption, alters placental structure and function, and reprograms offspring growth and liver transcriptomes in a sex-specific manner. This work identifies the preconception window as a critical period of vulnerability for EDC impacts on reproductive success and intergenerational health.

## Introduction

Infertility, defined as the inability to conceive after 12 months of unprotected intercourse, affects approximately 10–15% of couples worldwide and is a growing public health concern with far-reaching social and economic consequences.^1–3^ Although infertility arises from diverse causes, including anatomical, genetic, and endocrine factors, a mounting body of evidence points to environmental chemical exposures as a significant, yet underappreciated contributor.^4–11^ In particular, endocrine-disrupting chemicals (EDCs), a class of synthetic compounds that interfere with hormonal signaling, have emerged as key suspects in the global decline in fertility rates.^12–19^

More than 85,000 synthetic chemicals are registered for commercial use in the United States, and over 1,400 have been identified as EDCs based on their ability to disrupt endogenous hormone activity.^20,21^ These chemicals are pervasive in modern life, found in plastics, personal care products, food packaging, pesticides, flame retardants, and household goods.^22,23^ While some EDCs have been banned or regulated, many remain in circulation, and humans are routinely exposed to complex mixtures of low-dose chemicals throughout their lifetimes.^24,25^ Epidemiological studies have associated EDC exposure with menstrual irregularities, anovulation, recurrent pregnancy loss, and adverse birth outcomes.^26–29^ However, establishing causality and identifying the precise biological mechanisms by which these exposures impair fertility has been hindered by the complexity of real-world exposures, ethical constraints in human studies, and limitations of traditional toxicology models that often focus on single compounds at high doses.

Phthalates are among the most ubiquitous and extensively studied EDCs with known effects on female reproductive health. These multifunctional chemicals are used as plasticizers in polyvinyl chloride (PVC) products, as well as solvents and fragrance carriers in cosmetics, lotions, perfumes, and other personal care items.^30–33^ Due to their non-covalent incorporation into consumer goods, phthalates readily leach into the environment, leading to chronic, low-level exposure via ingestion, inhalation, and dermal absorption.^34^ Biomonitoring studies have consistently detected phthalate metabolites in human urine, blood, and even follicular fluid, with women often showing higher body burdens than men, likely due to gendered differences in product use.^35^ Experimental research has demonstrated that phthalate exposure during pregnancy can impair ovarian steroidogenesis, alter uterine gene expression, reduce endometrial receptivity, and increase the risk of implantation failure, pregnancy loss, and obstetric complications such as preeclampsia and preterm birth.^35–44^ However, little is known about preconception exposure or the mechanisms through which it affects fertility, fecundity, and offspring health outcomes.

The preconception period represents a particularly vulnerable window during which EDCs can disrupt key physiological processes required for successful reproduction. Beginning at puberty, female reproductive hormones undergo tightly regulated cyclical fluctuations that orchestrate ovulation and prepare the uterus for implantation.^45,46^ While this hormone-driven process is intentionally disrupted by contraceptives to prevent pregnancy, widespread exposure to environmental EDCs, found in plastic packaging, cosmetics, and feminine hygiene products, may unintentionally interfere with the same endocrine pathways.^12,47–50^ These exposures are continuous, low-dose, and largely unavoidable, making their effects on reproductive readiness especially concerning. Perturbations to hormonal signaling during the preconception window, defined here as the period between puberty and pregnancy, could impair oocyte maturation, ovulation, uterine receptivity, and implantation. Although a growing number of epidemiological studies have reported associations between EDC exposure and reduced fertility, establishing causality and elucidating mechanisms remains challenging.^51–54^ To overcome these barriers, the current study leverages a murine model to investigate how preconception exposure to an environmentally relevant phthalate mixture affects female reproductive physiology, implantation success, placenta form and function, and offspring health outcomes. We hypothesize that preconception phthalate exposure will disrupt hormone dependent processes including the estrus cycle and implantation, leading to downstream consequences for placental architecture and function and offspring health. By leveraging a physiologically relevant exposure paradigm and focusing on an etiologically important window of reproductive vulnerability, this work aims to elucidate the causal links between EDCs and reproductive success, with implications for understanding infertility, pregnancy loss, and intergenerational health effects in exposed populations.

## Materials and Methods

### Animals

All animal care, maintenance, and experimental procedures adhered to the standards outlined by the Animal Welfare Act and the U.S. Department of Health and Human Services Guide for the Care and Use of Laboratory Animals and were approved by the Clemson University Institutional Animal Care and Use Committee (IACUC). The Animal Research: Reporting of *In Vivo* Experiments (ARRIVE) Guidelines Checklist for Reporting Animal Research was used in construction of this manuscript with all elements met.^55^ A supervising veterinarian oversaw and monitored all procedures throughout the duration of the study. For each aim, female and male CD-1 mice were obtained from Charles River Laboratories (Raleigh, North Carolina) and housed at the Godley-Snell Research Center at Clemson University, an AAALAC-accredited biological resource facility. Animals were maintained under standard laboratory conditions with a controlled temperature of 25°C, 12:12 hour light-dark cycle, and average humidity between 45% and 60%. Mice were allowed a two-week acclimation period prior to initiation of experimental procedures. Housing and husbandry practices were specifically designed to minimize unintended exposure to EDCs. All mice were provided a soy-free diet (Teklad 2020), housed in poly-sulfone cages, supplied filtered drinking water via glass water bottles fitted with metal sippers, and bedded with woodchip material, in accordance with best practice guidelines for endocrine disruptor research.^56–61^

### Dosing prep

Oral exposure to a phthalate-mixture dissolved in a corn oil vehicle was used as the route of administration for all animals. To prepare the concentrated stock solution, individual phthalates were pipetted directly into a pre-weighed vial placed on an analytical balance. The volume of each chemical added was recorded to precisely achieve the target mass, based on the relative percentage composition calculated from previous assessments of human exposure.^62–64^(**Table 1**). The final mixture included benzylbutyl phthalate (BzBP; CAS No. 85-68-7), diisobutyl phthalate (DiBP; CAS No. 84-69-5), diisononyl phthalate (DiNP; CAS No. 28553-12-0), dibutyl phthalate (DBP; CAS No. 84-74-2), di(2-ethylhexyl) phthalate (DEHP; CAS No. 28553-12-0), and diethyl phthalate (DEP; CAS No. 84-66-2). Phthalates were obtained from Sigma-Aldrich (purity: ≥98%). Tocopherol-stripped corn oil (MP Biomedicals) served as the vehicle control for all dosing groups (**Fig.1**). The proportional composition of the mixture was based on urinary phthalate metabolite concentrations measured in pregnant women enrolled in the Children’s Environmental Health Research Center at the University of Illinois.^64–66^ Metabolite concentrations were used to recalculate the proportions of diester phthalates representing typical human exposures (**Table 1**). Working dosing solutions were prepared to achieve the targeted exposure level of 200 μg/kg/day by serial dilution of the concentrated stock in corn oil. All dosing solutions were mixed overnight on an orbital shaker and stored in amber glass vials to prevent photodegradation until use.

**Figure 1.**
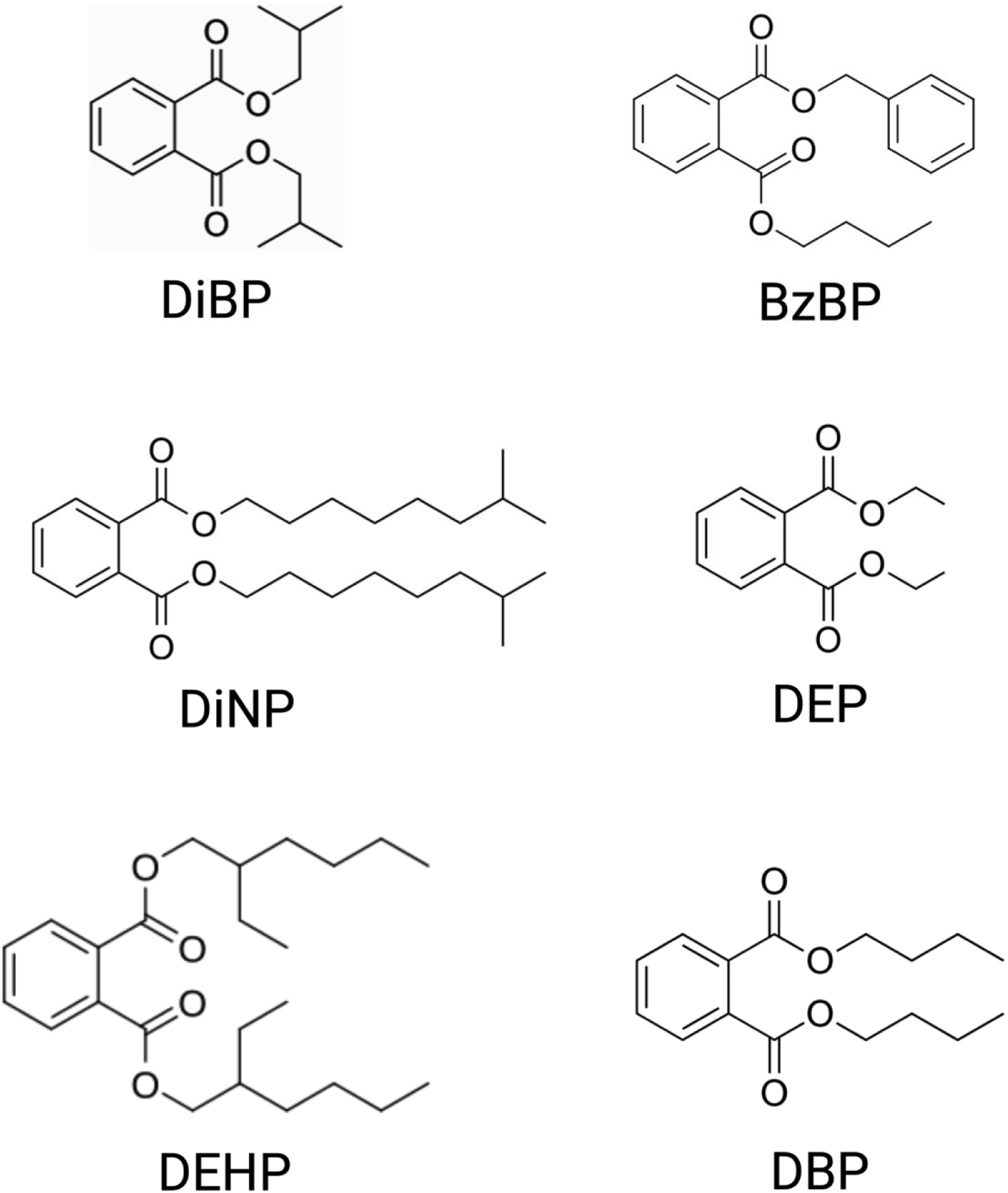
Chemical structures of the six phthalates that make up the phthalate mixture in this study. Diisobutyl phthalate (DiBP), diisononyl phthalate (DiNP), di(2-ethylhexyl) phthalate (DEHP), benzylbutyl phthalate (BzBP), diethyl phthalate (DEP), and dibutyl phthalate (DBP).

**Table 1.** Phthalates used in mixture, percentage of each phthalate in mixture, and calculated daily exposure in mice with relevance to human daily exposures.

| Phthalate | Abbreviation | % of Dosing Mixture | Mouse Exposure (µg/kg/day) * | Human Exposure (µg/kg/day)<sup>65,66</sup> |
| --- | --- | --- | --- | --- |
| Diisobutyl Phthalate | **DiBP** | 8 | 1.6 – 16 | 0.12 – 1.4 |
| Diisononyl Phthalate | **DiNP** | 15 | 3 – 30 | ≤ 26 |
| Di(2-ethylhexyl) Phthalate | **DEHP** | 21 | 4.2 – 42 | 3 – 30 |
| Benzylbutyl Phthalate | **BzBP** | 5 | 1 – 10 | 0.26 – 0.88 |
| Diethyl Phthalate | **DEP** | 35 | 7 – 70 | 2.3 – 12 |
| Dibutyl Phthalate | **DBP** | 15 | 3 – 30 | 0.84 – 5.22 |

\*Mouse exposure based on the selected 200µg/kg/day dose.

### Animal Husbandry and Exposure

Adult CD-1 mice (8 females, 4 males) were obtained from Charles River Laboratories (Raleigh, NC). Breeding pairs were established by housing two females with a single male to generate a colony of in-house offspring for exposure. To ensure uniformity in pubertal development at the start of exposure, female offspring were weaned at PND 21 and randomly selected for experimental use at postnatal day (PND) 30. All animals were weighed daily throughout the exposure period, and estrous cyclicity was tracked by daily vaginal smear (∼10:00 AM) using sterile physiological saline (0.9% NaCl) as described previously.^67^ Lavage samples were stained using 0.05% toluidine blue to identify estrous cycle stages, serving as non-invasive biomarkers of endocrine and reproductive function (**Supplemental Fig. 1**).

Beginning on PND 30, female mice were randomly assigned to receive either the 200 μg/kg/day phthalate-mixture (n = 25) or vehicle control (n = 20) via daily oral pipetting for 30 consecutive days, concluding at PND 60. All doses were given in 21 – 25 μl volumes based on their body weight. Following the final day of dosing at PND 61, 2 virgin female mice were paired with a single non-littermate male to maintain outbred genetic heterogeneity. Upon detection of a vaginal sperm plug, males were removed, and dams were singly housed. The morning a sperm plug was detected was considered embryonic day (E) 0.5. Pregnant dams were randomly assigned to one of three cohorts defined by their experimental endpoint, cohort 1 to assess implantation, cohort 2 to assess placental structure and function, and cohort 3 to evaluate offspring reproductive and metabolic outcomes (**Fig. 2**).

**Figure 2:**
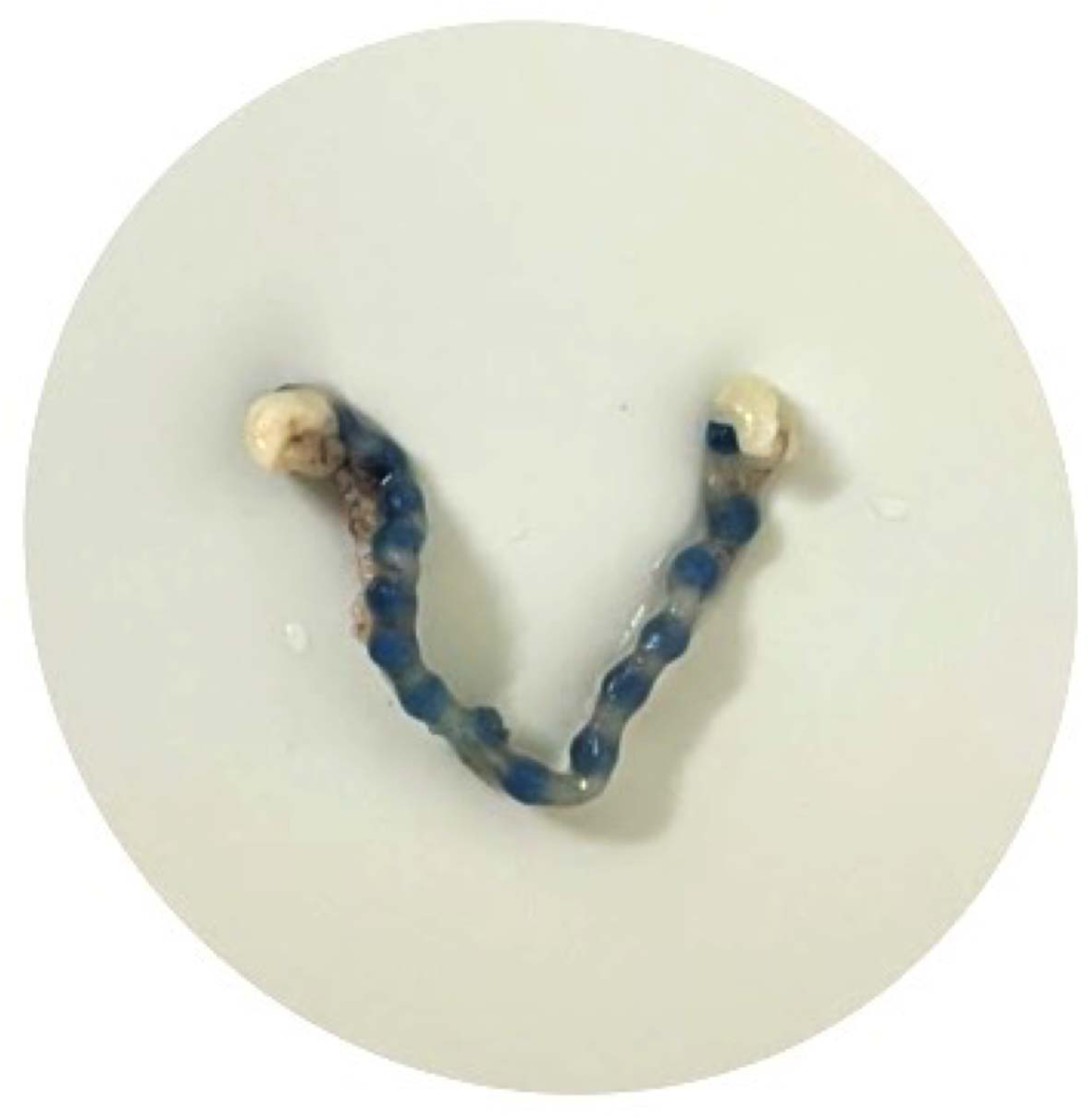
Representative image of Chicago blue stained implantation sites.

### Tissue Collection

Animals were sacrificed using a method approved by the Panel on Euthanasia of the American Veterinary Medical Association. For cohort 1, mice were fully anesthetized with isoflurane inhalant prior to tail vein dye injection. Cohorts 2 and 3 were euthanized via carbon dioxide (CO_₂_) asphyxiation immediately followed by rapid decapitation.

*Cohort 1:* Implantation success was assessed on E5 by intravenous tail vein injection of 1% Chicago Blue dye (Sigma-Aldrich) in sterile saline for dams exposed to 200 μg/kg/day phthalate-mixture (n = 4) or corn oil (n = 3). Uteri were subsequently examined for dye localization marking implantation sites (**Fig. 3**).

**Figure 3:**
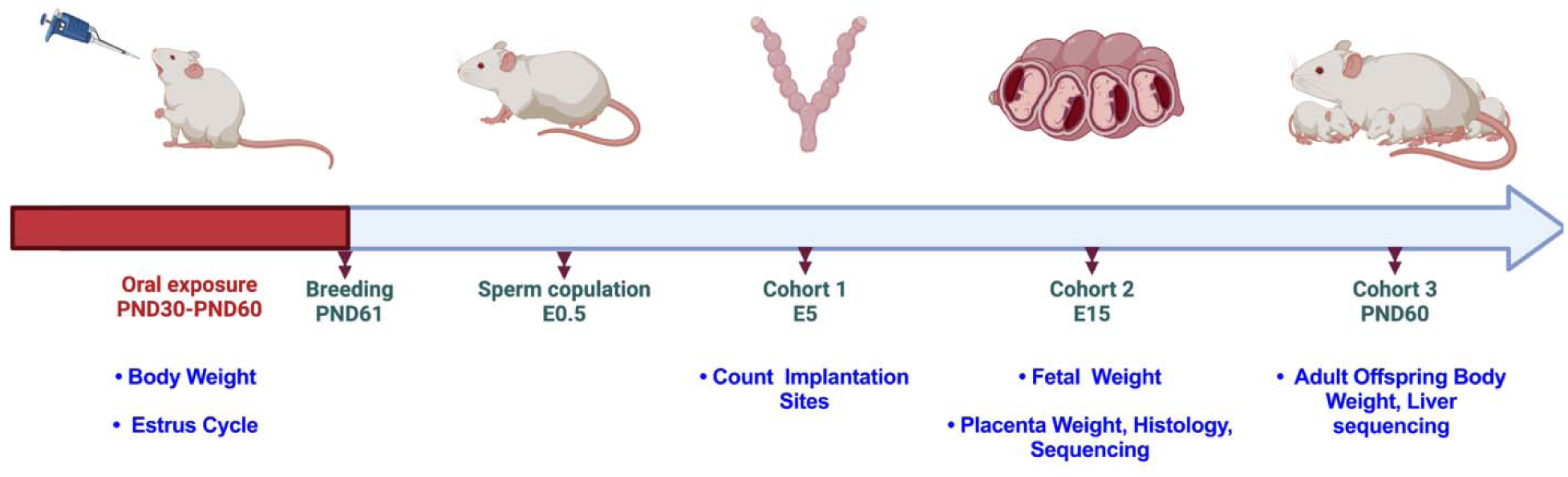
Timeline of experimental design. Pregnant females were split into three cohorts to evaluate impacts on: (1) implantation (E5), (2) fetal and placental development (E14.5), and (3) adult offspring health outcomes (PND 60). Maternal body weight and estrus cycle were tracked throughout the duration of exposure. Implantation sites were counted at E5. Litter size was counted, and fetal and placental weights were collected at E14.5. Adult offspring bodyweight and estrus cycle were tracked between PND 30-60. Finally, tissues including livers, placentas were collected for RNA-sequencing.

*Cohort 2:* Fetuses and placentas were collected at E14.5 from 200 μg/kg/day phthalate-mixture (n = 5) and control dams (n = 4). Tissues were weighed immediately after collection. A small portion of tail tissue was collected from each fetus for sex determination by genotyping using primers specific to Jarid1 (5’-TGAAGCTTTTGGCTTTGAG-3’ and 5’- CCGCTGCCAAATTCTTTGG-3’) as described elsewhere.^68^ Each placenta was bisected, with one half flash-frozen for RNA sequencing and stored at –80°C, and the other half fixed in 10% neutral buffered formalin (NBF) for histopathological analysis. Based on the results of sex genotyping, one male and one female from each litter were selected for downstream analyses.

*Cohort 3:* At weaning, dams were sacrificed, and livers were collected and flash-frozen for downstream analysis (200 μg/kg/day n = 3, control n = 3). Postnatal weight change for two adult offspring per sex following maternal exposure per sex was monitored from PND30 to 60 weekly (200 μg/kg/day n = 15 per sex, control n = 9 male and n=11 female). Females born in this group were monitored daily to assess estrous cycle from PND 30 to 45 as well. Adult offspring livers and gonads were collected on PND 60 for both sexes and flash-frozen for downstream analysis (200 μg/kg/day n = 3, control n = 3).

### Histology

For gross histopathological assessment, a half of a placenta, 1 per sex per litter, was initially fixed in 10% neutral buffered formalin (NBF). Tissues were then sequentially transferred to NBF containing 30% sucrose, followed by immersion in a solution of 30% sucrose in phosphate-buffered saline (PBS) for cryoprotection. Fixed placentas were stored at –80_°C until histological processing. Prior to paraffin embedding, tissues underwent a reverse sucrose gradient protocol, 20% for 24 hours, 10% for 24 hours, and PBS for 24 hours, to remove sucrose prior to paraffin processing. Placentas from control dams (n = 4/sex) and 200 μg/kg/day phthalate mixture group (n = 5/sex) were sent to the Clemson Light Imaging Facility (CLIF) for hematoxylin and eosin (H&E) staining and imaging. Tissues were embedded in paraffin using the Leica Histocore Arcadia Embedding Station, sectioned at 5_μm thickness with the Leica Histocore Biocut Rotary Microtome, and mounted on glass slides. Sections were de-waxed and rehydrated using a series of xylene and ethanol baths, where ethanol concentrations progressively decreased from 100% - 50% followed by a final rinse in PBS. Slides were stained with hematoxylin and eosin (H&E) for morphological analysis using an H&E staining kit (Abcam) according to the manufacturers protocol. Post-processing and staining, two female placentas from the control group were excluded from the study due to compromised tissue integrity (n = 2 female control placentas). Images were taken using a DSX1000 Digital Microscope (Olympus). Area measurements for the whole placenta and specific placental zones (labyrinth, junctional and decidua) were determined using ImageJ (version1.54g). Areas were measured by two blinded researchers and the average area from those two were used for data analysis.

### RNA-Sequencing

RNA-seq as an initial hypothesis-generating approach was performed on E14.5 placentas and PND 60 offspring. Approximately 30mg of tissues were used for RNA extraction with the Qiagen RNEasy Miniprep kit according to the manufacturer protocol (Qiagen, Cat. 74134). Total RNA was submitted to Novogene (Sacramento, CA, USA) for mRNA sequencing using the Illumina platform. RNA quality was determined using an Agilent 5400 Bioanalyzer and only samples that passed Novogene quality control criteria were used for sequencing. Placenta (control n = 4/sex and 200 μg/kg/day phthalate mixture n = 5/sex), offspring liver (control female n = 3, control male n = 2, 200 μg/kg/day phthalate mixture n = 3/sex), and offspring gonad (control n = 3/sex and 200 μg/kg/day phthalate mixture n = 3/sex) messenger RNA was isolated from total RNA using poly-T oligo-attached magnetic beads, followed by heat fragmentation and first-strand cDNA synthesis using random hexamer primers. Second-strand synthesis was performed using dUTP to maintain strand specificity.^69^ Libraries underwent end repair, A-tailing, adapter ligation, size selection, PCR amplification, and final purification. Library quality and concentration were assessed using Qubit fluorometry, real-time PCR, and an Agilent 5400 Bioanalyzer. Quantified libraries were be pooled and sequenced on Illumina platforms, according to effective library concentration and data amount.

### RNA-seq Data Processing

#### Data Quality Control

Raw data (raw reads) of fastq format were firstly processed through in-house perl scripts. In this step, clean data (clean reads) were obtained by removing reads containing adapter, reads containing ploy-N and low-quality reads from raw data. At the same time, Q20, Q30 and GC content the clean data were calculated. All the downstream analyses were based on the clean data with high quality. The transcriptome data discussed in this publication has been deposited in NCBI’s Gene Expression Omnibus (PRJNA1308639).

#### Read mapping to the reference genome

Reference genome and gene model annotation files were downloaded from genome website directly. Index of the reference genome was built using Hisat2 v2.0.5 and paired-end clean 1 reads were aligned to the reference genome using Hisat2 v2.0.5. We selected Hisat2 ^70^as the mapping tool for that Hisat2 can generate a database of splice junctions based on the gene model annotation file and thus a better mapping result than other non-splice mapping tools.

#### Quantification of gene expression level

Feature Counts ^71^ v1.5.0-p3 was used to count the reads numbers mapped to each gene. Average uniquely mapped reads in placentas, livers, and gonads ranged between 37 - 38 million per library. And then FPKM of each gene was calculated based on the length of the gene and reads count mapped to this gene. FPKM, expected number of Fragments Per Kilobase of transcript sequence per Millions base pairs sequenced, considers the effect of sequencing depth and gene length for the reads count at the same time, and is currently the most used method for estimating gene expression levels.

#### Differential expression analysis

Differential expression ^72,73^ analysis of two conditions/groups (two biological replicates per condition) was performed using the DESeq2Rpackage (1.20.0). DESeq2 provide statistical routines for determining differential expression in digital gene expression data using a model based on the negative binomial distribution. The resulting P-values were adjusted using the Benjamini and Hochberg’s approach for controlling the false discovery rate. Genes with an adjusted P-value ≤ 0.05found by DESeq2 were assigned as differentially expressed. For edgeR without biological replicates: Prior to differential gene expression analysis, for each sequencing library, read counts were 4 adjusted using the edgeR R package (4.0.16) by scaling normalization factors to eliminate differences in sequencing depth between samples, followed by differential expression analysis^74^. The resulting P value is adjusted using the Benjamini and Hochberg’s methods to control the error discovery rate. The threshold of significant differential expression: padj <= 0.05 & |log2(foldchange)| >= 1. Adult offspring liver data was further filtered due to the low sample size for males according to previously published protocols.^75^ Briefly, data was filtered to exclude transcripts that were detected in < 2 samples/group, which brought the total number of genes for male livers to 14,896 and 14,078 for females.

#### Statistical Methods

Statistical analysis was performed using GraphPad Prism version 10 (version 10.4.2, La Jolla, California) and the R statistical environment (version 4.4.3, R core team 2021) with statistical significance set at *alpha* = 0.05. Results are presented as mean ± SEM or min to max. For comparisons between two groups, unpaired Student’s *t*-tests were used. Two-way ANOVA followed by LSD post hoc tests was applied for multiple group comparisons. For t-test’s and ANOVA’s effect size was determined by calculating an Eta squared (η^2^), effects of which are defined as small at 0.01, medium at 0.06, and large at 0.14.^76^ For sequencing results over- representation analysis (ORA) was performed using the enrichGO() and enrichKEGG() functions on the subset of significantly differentially expressed genes. Enrichment scores were calculated as −log10(adjusted p-value), with significance set at adjusted p-values (FDR) < 0.05. These functions identify Gene Ontology (GO) terms and Kyoto Encyclopedia of Genes and Genomes (KEGG) pathways that are statistically over-represented, irrespective of whether they were up or downregulated among the DEGs.

## Results

### Maternal bodyweight, estrus cycle, and fertility

Maternal body weight was tracked from PND 30 to 60. No significant differences in weight gain over time or total weight gained were observed in exposed dams compared to controls (**Fig. 4A and 4B**). Maternal estrus cycle appeared to impacted by phthalate exposure, with more time spent in proestrus (F (24, 19) = 2.258, *p* = 0.07, η^2^ = 0.07) and less time spent in metestrus (F (24, 19) = 2.283, *p* = 0.09, η^2^ = 0.06), but the effect did not reach statistical significance (**Fig. 4C**). Phthalate exposure also had no significant effect on the number of implantation sites at E5 (**Fig. 4D**) or the number of pups per litter at E 14.5 (**Fig. 4E**). Collectively, tracking maternal health during the exposure period provided suggestive evidence of endocrine disruption, as indicated by altered estrous cycling in phthalate-exposed dams.

**Figure 4.**
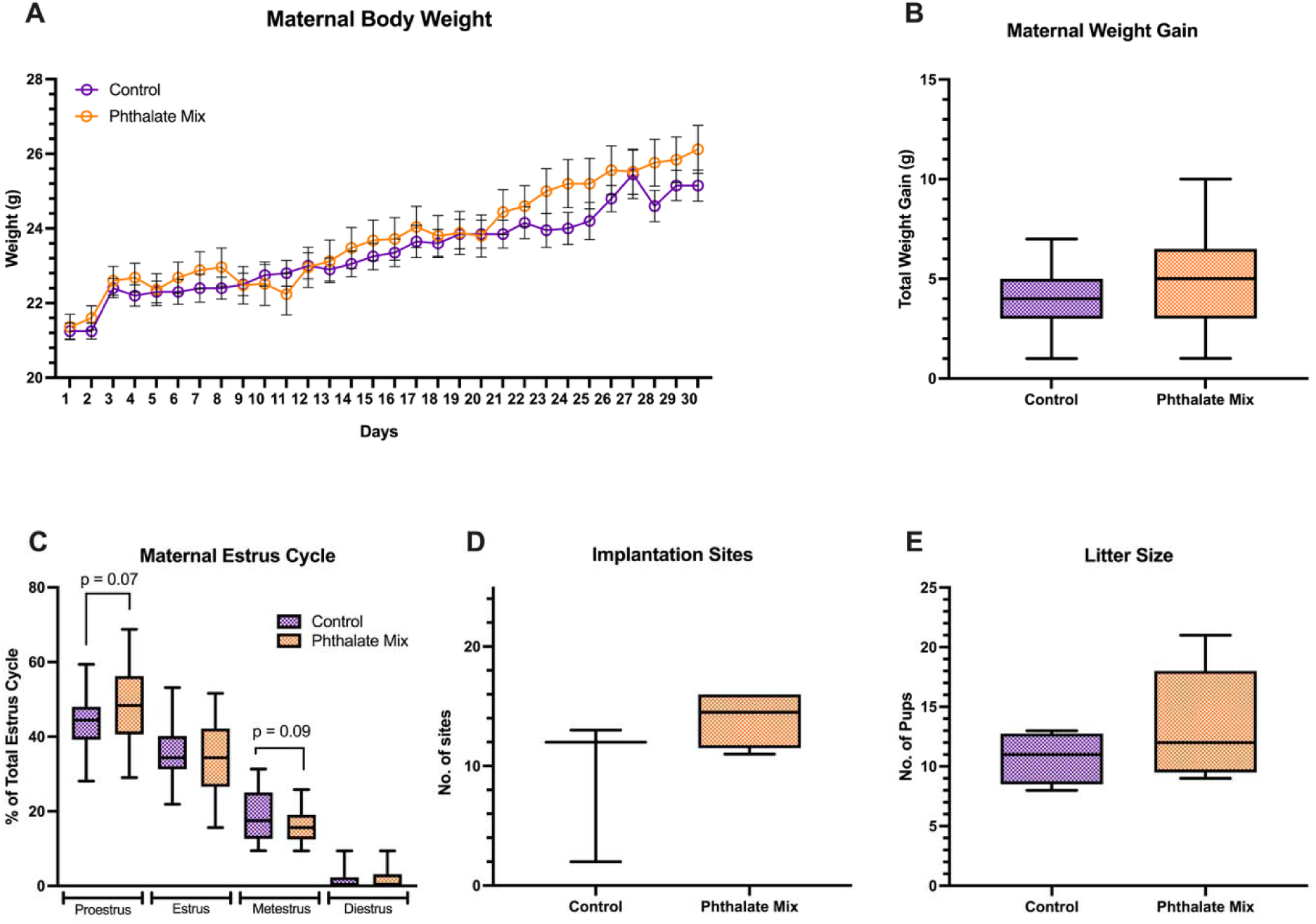
Dams exposed to phthalates during the preconception window exhibited no significant alterations in body weight or fertility, though modest disruptions in estrous cycling were observed. Phthalate exposure had no significant impact on maternal bodyweight (A) over time or (B) total weight gain. No significant effect of exposure was observed on (C) number of days in each stage of the estrus cycle; however, t-test revealed a suggestive increase in the number of days in proestrus and decrease in metestrus compared to controls; n=20 control dams; n=25 phthalate Mix dams. Phthalate exposure didn’t significantly change the number of (D) implantation; n=3 control dams’ size; n=4 PM litter size or (E) pups per litter; n=4 control dams; n=5 phthalate Mix dams. Line graph depicts mean ± SEM. Boxplots depict mean, min, and max. Line graphs depict mean ± SEM.

### Offspring bodyweight, placental weight, and placental histology

We next evaluated the impact of phthalate-mixture exposure on pregnancy outcomes by assessing changes in fetal and placental development at E14.5. One litter from the phthalate exposed group was removed from the analysis due to having an abnormally large litter size leading to overcrowding in the uterus. Exposure significantly increased average fetal bodyweight relative to control group at E14.5 (F (3, 3) = 5.237, *p* = 0.02, η^2^ = 0.64; **Fig. 5A**). However, phthalate treatment did not significantly impact average placental weight (**Fig. 5B**) or Fetal:Placental (F:P) weight ratio (estimate of placental efficiency) compared to the control group (**Fig. 5C**). Exposure also had no significant effect on the number of resorptions, litter size, or the distribution of pups between the left and right uterine horns at E14.5 (**Supplemental Data 1, Table S6**).

**Figure 5.**
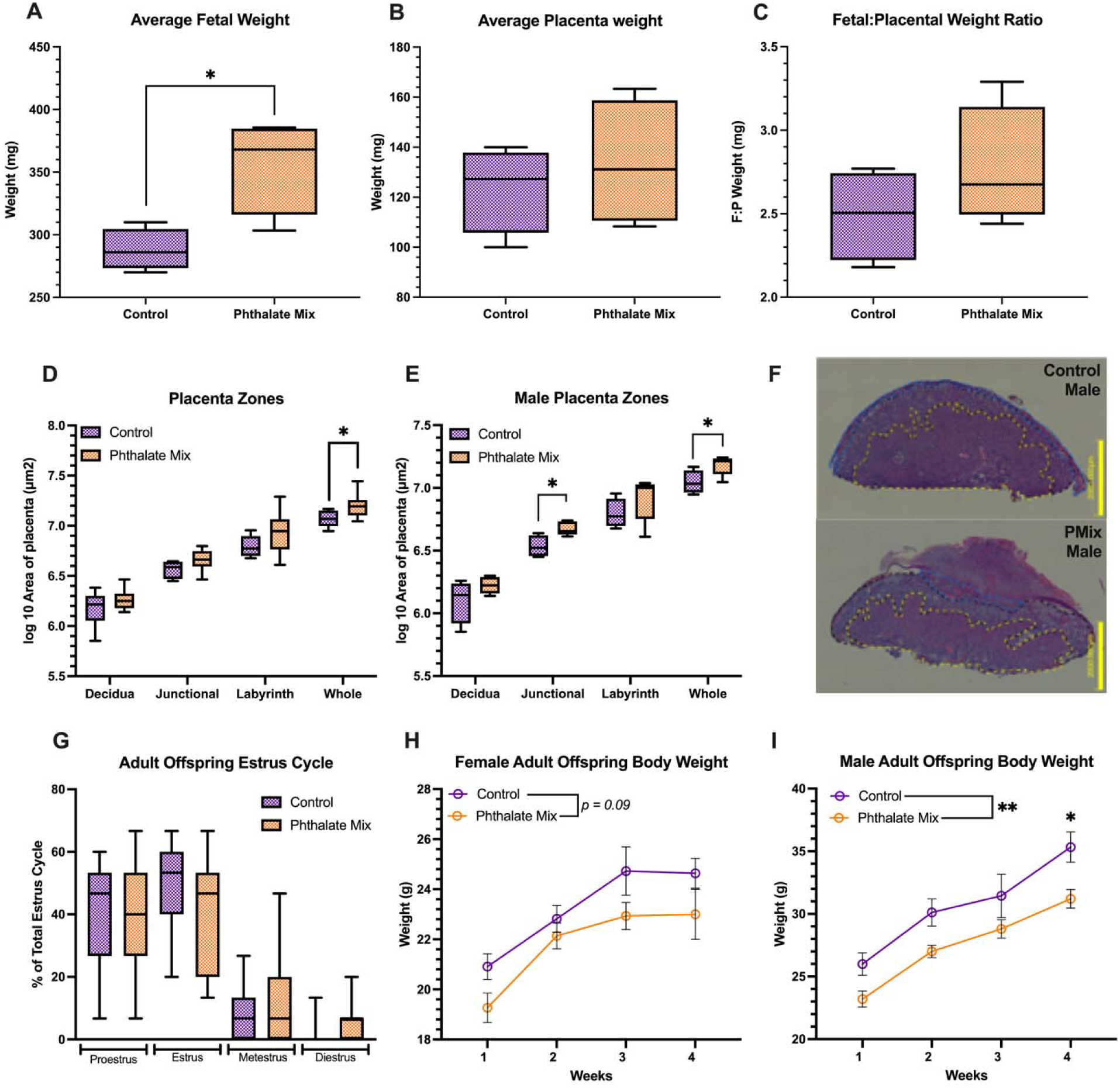
Preconception phthalate exposure significantly influenced offspring body weight during both gestation and adulthood and exerted sex-specific effects on placental morphology, most notably an increase in the male junctional zone. (A) Exposure significantly increased fetal body weight but had no effect on (B) placental weight or (C) fetal to placental (F:P) weight ratio, (an estimate of placental efficiency). n=4 control dams’ fetuses; n=5 phthalate Mix dams’ fetuses. (D) A significant effect of exposure on placental areas was observed in whole placenta with (E) Male offspring of exposed dams had a significantly larger junctional zone and whole placenta area; n *=* 2 female control; n *=* 4 male control; n *=* 5 per sex per exposed group. (F) Representative male placental histology from control and exposed groups. (G) Adult offspring estrus cycle from PND30-45 showed no significant difference between control and phthalate mix group. Adult offspring (H) female body weight did not significantly differ between treatment groups, but (I) male offspring of exposed dams had significantly lower body weight; n *=* 11 female control; n *=* 9 male control; n *=1* 5 female phthalate Mix and n=15 male phthalate Mix. Boxplots depict mean, min, and max. Line graphs depict mean ± SEM. Line graphs depict mean ± SEM.; \**p* ≤ 0.05; \*\**p* ≤ 0.01

Although the overall placental architecture appeared intact, gross pathological examination suggested subtle alterations associated with phthalate exposure. To visualize structural differences across placental regions, histological sections from both sexes of control and exposed groups were examined, with the decidua, junctional zone, and labyrinth distinctly separated and analyzed. To meet the assumptions of parametric testing, area measurements of each placental zone were log₁₀-transformed to ensure normality and stabilize variance across groups.^77^ Exposure significantly increased the size of the whole placenta (F (9, 5) = 2.106, *p* = 0.04, η^2^ = 0.27; **Fig. 5D**). When stratified by sex, a significant increase in the junctional zone (F (3, 4) = 2.524, *p* = 0.02, η^2^ = 0.57) and whole placenta (F (3, 4) = 1.359, *p* = 0.05, η^2^ = 0.44) area was observed in male placentas of the exposed group compared to controls (**Fig .5E and 5F**). However, given the limited sample size in the control female group (n = 2), the study was underpowered to robustly detect sex-specific differences. These findings suggest a region-specific and potentially sex-dependent sensitivity of the placenta to preconception phthalate mixture exposure, specifically implicating the junctional zone.

Estrus cycle and bodyweight were also evaluated for adult offspring. No significant effect of exposure was observed for adult female offspring estrus cycle (**Fig. 5G**). However, maternal phthalate exposure significantly altered offspring body weight. While not significant, exposure had a suggestive impact on adult female offspring bodyweight (F (1, 24) = 2.970*, p* = 0.09, η^2^ = 0.11; **Fig. 5H**). A significant effect of phthalate exposure on adult male offspring bodyweight was observed (F (1, 22) = 8.243, *p* = 0.009, η^2^ = 0.27; **Fig. 6I**)

**Figure 6.**
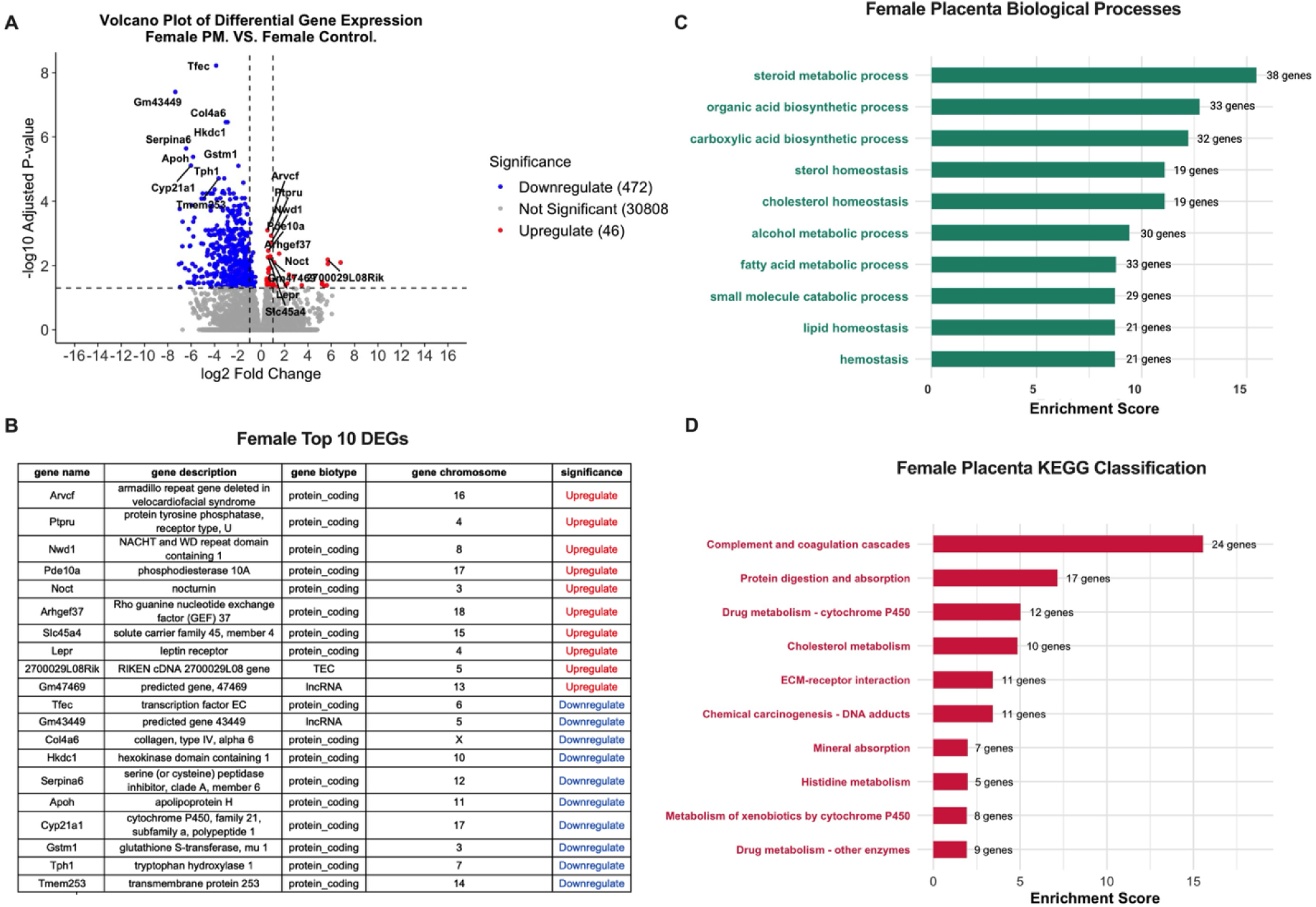
Preconception phthalate exposure markedly altered the female placental transcriptome, with differential expression of genes involved in steroid metabolism, sterol homeostasis, and xenobiotic processing. (A) Volcano plot demonstrating upregulation in 472 genes and down regulation in 46 genes following preconception exposure to phthalate mix (B) list of top 10 genes upregulated and down regulated (C) GO analysis on biological processes suggested changes in lipid and steroid metabolic processes (D) KEGG pathway analysis noted alternations in xenobiotic metabolism, steroid biosynthesis, and extracellular matrix organization

### Placenta – DEGs, Gene Ontology (GO) enrichment, and Kyoto Encyclopedia of Genes and Genomes (KEGG) overview in placenta

Principal components analysis (PCA) showed little separation between male and female control and exposed placentas (**Supplemental Fig. 2B**). However, a heatmap of all the differentially expressed genes showed cleared separation via hierarchical clustering of control and exposed placentas (**Supplemental Fig. 2C**). Differential gene expression analysis (contrast matrix exposed female – control female or exposed male – control male; *p* value ≤ 0.05; log2FoldChange ≥ 0) revealed significant changes in individual gene expression (p-adj ≤ 0.05). A total of 518 differentially expressed genes (DEGs) were identified in female placenta, including 472 downregulated and 46 upregulated genes (**Supplemental Table S9**). However, in male placenta exposure only induced a significant upregulation in 9 genes. Collectively these findings suggest that exposure had a greater impact on the female placental transcriptome compared to males.

In the female placenta, majority of the significant differentially expressed genes were downregulated (**Fig. 6A**). Notable genes among the top 10 upregulated and downregulated genes include *Slc45a4* (sugar transporter), *Lepr* (regulates growth and nutrient transport), *Cyp21a1* (synthesis of steroid hormones), and *Tph1* (biosynthesis of serotonin; **Fig. 6B**). To capture functional patterns among differentially expressed genes, Gene Ontology (GO) enrichment. Significant enrichment for biological processes in females included steroid metabolic processes, cholesterol homeostasis, and lipid homeostasis (**Fig. 6C**). KEGG pathway analysis provided additional mechanistic insights, focusing on canonical signaling and metabolic routes impacted by differential gene expression. The most significantly enriched pathway in female placenta was Complement and coagulation cascades (24 genes), followed by protein digestion and absorption (17 genes) and drug metabolism-cytochrome P450 (12 genes). (**Fig. 6D**).

In the male placenta, all 9 significant differentially expressed genes were upregulated (**Fig. 7A**). Majority of these genes encode for biological molecules involved in gene regulation, including long non-coding RNAs and transcription factors (**Fig. 7B**). Significant enrichment of biological processes such as RNA splicing, negative regulation of gene expression, and regulation of mRNA processing were observed. Given the small number of significantly enriched genes, only a single KEGG classification pathway was identified, cytoskeleton in muscle cells.

**Figure 7:**
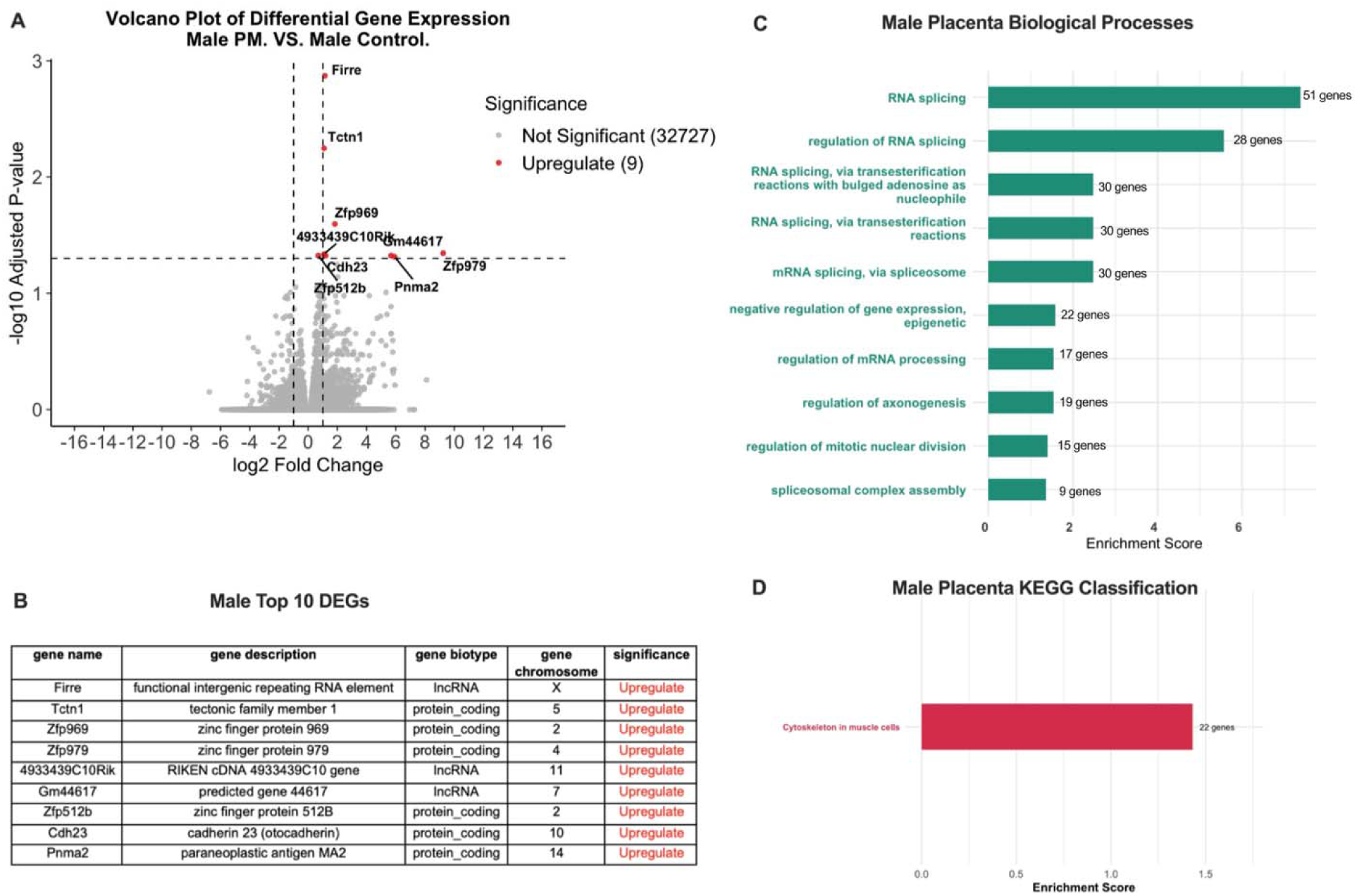
Preconception phthalate exposure produced modest transcriptomic changes in the male placenta, primarily affecting genes associated with RNA processing. (A) Upregulation in 9 genes following dams’ exposure to phthalate mix (B) list of top 10 genes upregulated and down regulated (C) GO analysis on biological processes suggested changes in RNA processing (D) only one pathway noted in KEGG analysis.

### Liver – DEGs, Gene Ontology (GO) enrichment, and Kyoto Encyclopedia of Genes and Genomes (KEGG) overview in placenta

PCA showed clear separation between the male and female liver transcriptome (**Supplemental Fig. 3B**). Furthermore, hierarchical clustering suggests that preconception phthalate exposure feminized offspring male livers and masculinized female livers with control females clustering more closely with exposed males and exposed females clustering with control males (**Supplemental Fig. 3C**). A total of 175 DEGs were identified in the female liver and 95 in the male liver (**Supplemental Tables S16 and S17**).

Similar to the placenta, significant DEGs in adult female offspring livers were predominantly downregulated (**Fig. 8A**). Among the top 10 up- and down-regulated genes were *Cyp4a10* (fatty acid metabolism), *Cyp7a1* (bile acid synthesis), *Mthfr* (folate metabolism), and *H2-DMB1* (antigen presentation and processing), among others (**Fig. 8B**). Significant enrichment of GO biological processes organic hydroxy compound transport, positive regulation of T cell activation, and organic acid biosynthetic processes was observed (**Fig. 8C**), however no significant KEGG classifications were identified. In the male liver, DEGs were predominantly upregulated (**Fig. 8D**). Genes including *Elf4* (cell differentiation and immune function), *Tnfrsf22* (cytokine signaling), *Adm2* (peptide hormone), and *Ly6a* (immune response) are among the top 10 up- and down-regulated genes in the male liver (**Fig. 8D**). No significant enrichment of GO biological processes or KEGG classifications were identified.

**Figure 8:**
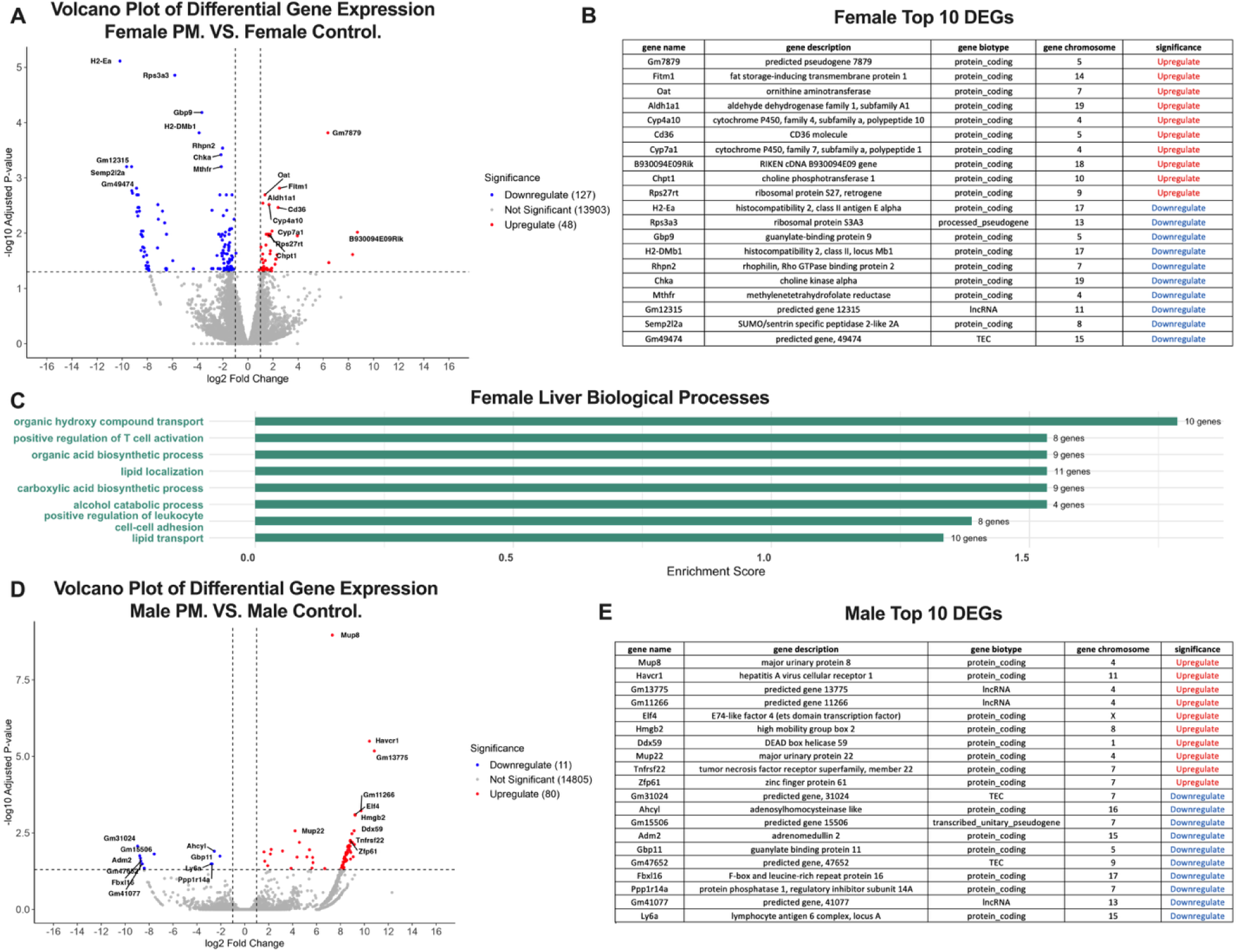
Significant transcriptomic alterations were observed in the adult offspring liver following preconception phthalate exposure, with enrichment of energy production– related genes in females. (A) Upregulation in 48 and downregulation in 127 following dams’ exposure to phthalate mix in female adult offspring (B) list of top 10 genes upregulated and down regulated (C) GO analysis on biological processes suggested changes in organic hydroxy compound transport, positive regulation of T cell activation, and organic acid biosynthetic processes (D) Upregulation in 80 and downregulation in 11 following dams’ exposure to phthalate mix in male adult offspring <u>(E)</u> list of top 10 genes upregulated and down regulated in male adult offspring liver.

## Discussion

Endocrine-disrupting chemicals (EDCs), including phthalates, are pervasive in the environment and are found in a wide array of consumer products such as plastics, cosmetics, and personal care items.^78,79^ Women of reproductive age are especially vulnerable to these exposures, which have been increasingly linked to reproductive dysfunction and adverse pregnancy outcomes. ^66,67^Although gestational exposure to phthalates has been extensively studied, emerging evidence suggests that exposures prior to conception may also disrupt critical early reproductive events and fetal programming. ^82–84^ However, the preconception window remains a poorly understood yet potentially sensitive period. This study provides new evidence that preconception exposure to a human-relevant phthalate mixture disrupts maternal endocrine function, alters placental structure and gene expression, and impairs offspring development in a sex-specific manner. By modeling environmentally relevant exposure during the preconception window, a critical yet understudied period, our findings expand current understanding of when and how EDCs may impact reproductive and developmental health.

### Preconception Exposure Alters Maternal Endocrine Function

Consistent with prior studies of phthalate-induced reproductive disruption ^85–88^, we observed that 30-day preconception exposure resulted in altered estrous cyclicity, specifically prolonged proestrus and shortened metestrus stages. Although no significant changes in maternal body weight or fertility outcomes (implantation or litter size) were detected, changes in cyclicity suggest that phthalates disrupted the hypothalamic-pituitary-gonadal (HPG) axis.^89^ These alterations occurred despite the absence of overt maternal toxicity, indicating a subtle but biologically meaningful endocrine effect during the reproductive prime. Disruptions to cyclicity may influence the timing and hormonal environment of conception, thereby indirectly affecting placental development and fetal health.^90–93^

### Placental Disruption and Fetal Growth Outcomes

While implantation rates and litter sizes were unaffected, we found that fetal body weight was significantly increased at E14.5 in exposed animals. This was accompanied by an expansion of the junctional zone in male placentas, suggesting altered trophoblast development or placental compensation in response to maternal stress. The junctional zone, which includes hormone-producing spongiotrophoblast cells, plays a central role in regulating fetal growth and maternal-fetal signaling. ^94^ Previous studies have shown that phthalate exposure during gestation can lead to abnormal placental morphology ^95–98^, and our results extend these findings to the preconception period.

Despite limited transcriptional changes in male placentas, these structural alterations were associated with postnatal consequences: male offspring exhibited significantly lower weight gain in adulthood, suggesting lasting metabolic or endocrine programming effects. These results support the concept that male placentas, while less transcriptionally plastic, may be more vulnerable to long-term physiological disruption.

### Female Placentas Exhibit Greater Transcriptional Sensitivity

Transcriptomic analysis revealed that female placentas were more transcriptionally responsive to preconception phthalate exposure, with over 500 differentially expressed genes compared to just 9 in male placentas. These sex-specific responses align with previous studies indicating that female placentas are more immunologically active and transcriptionally adaptable.^99–102^ Enrichment analyses pointed to disruptions in immune regulation (complement and coagulation cascades), xenobiotic metabolism, steroid biosynthesis, and extracellular matrix organization— processes critical to placental integrity and function.

Notably, we observed downregulation of *Serpina6*, which encodes corticosteroid-binding globulin (CBG)^103^, a regulator of glucocorticoid bioavailability. Disruption of glucocorticoid signaling may impair fetal stress responses and immune modulation.^104,105^ Additionally, *Gstm1*, a key antioxidant gene involved in glutathione-mediated detoxification^106^, was suppressed, suggesting increased oxidative stress vulnerability—a well-documented mechanism of phthalate toxicity.^107–109^

### Pathways Implicated in Placental Vascularization, ECM Remodeling, and Metabolism

Other key targets included *TFEC*, a transcription factor interacting with *TFEB* to regulate placental vascularization via VEGF signaling.^110,111^ Alterations in this pathway may explain impaired labyrinth zone formation and angiogenesis previously reported following DEHP exposure.^112^ Similarly, changes in ECM-related genes such as *Col4a6* and *Arvcf* point to disrupted trophoblast invasion and decidualization, hallmarks of healthy placentation.^113–115^ The observed suppression of these genes suggests impaired remodeling processes.

Our KEGG and GO analyses also implicated disruptions in lipid metabolism and steroid hormone biosynthesis. Upregulation of *Nocturnin* (NOCT), a circadian-regulated regulator of lipid handling, suggests altered placental energy homeostasis. NOCT has been linked to resistance to diet-induced obesity and impaired insulin signaling in mouse models, raising questions about its role in fetal metabolic programming.^116,117^ Dysregulation of placental lipid balance could impair steroidogenesis, nutrient transport, and fetal growth trajectories.^118^

### Potential Impacts on Neurodevelopment and Epigenetic Programming

We also identified changes in genes associated with neurodevelopment and epigenetic regulation. Downregulation of *Tph1*, a key enzyme in placental serotonin synthesis, may reduce fetal serotonin availability during critical neurodevelopmental windows^119–122^. Additionally, altered expression of multiple long non-coding RNAs (lncRNAs) supports growing evidence that phthalates can induce epigenetic reprogramming with long-term consequences^123–125^. These transcriptional changes may serve as early biomarkers of disrupted developmental trajectories.

Male placentas, though transcriptionally less responsive, showed enrichment of pathways related to RNA splicing and post-transcriptional regulation. Emerging literature suggests that EDCs such as BPA and phthalates can impair RNA-binding protein activity, disrupt splicing, and interfere with mRNA translation, potentially contributing to neurodevelopmental disorders (e.g., autism) even in the absence of strong transcriptional effects.^126,127^

### Adult Offspring Hepatic Disruption and Growth Outcomes

Preconception phthalate exposure elicited sex-specific transcriptomic changes in the adult offspring liver. PCA and hierarchical clustering revealed a “feminization” of male livers and “masculinization” of female livers, suggesting disruption of normal sex differences in hepatic gene expression. In females, upregulation of metabolic genes such as *Cyp4a10* and *Cyp7a1* indicates enhanced fatty acid oxidation and bile acid synthesis, processes that, when dysregulated, can disrupt lipid homeostasis and energy balance. While increased bile acid synthesis may initially support nutrient absorption, excessive activity may impair cholesterol regulation and contribute to altered metabolic programming. Concurrent downregulation of *Mthfr* points to disrupted one-carbon metabolism, with implications for epigenetic regulation and endocrine signaling. In males, several immune- and signaling-related genes were differentially expressed, with *Elf4* and *Tnfrsf22* upregulated but *Adm2* and *Ly6a* downregulated. Loss of *Adm2*, a peptide hormone involved in vascular tone and metabolic regulation, and *Ly6a*, a modulator of immune responsiveness, suggests impaired hepatic signaling capacity and reduced adaptability to stress. These changes could compromise the balance between immune regulation and growth hormone/IGF pathways, constraining anabolic growth. Together, these liver disruptions provide a mechanistic link to the reduced adult bodyweight observed in both sexes, statistically significant in males and trending downward in females, through sex-specific alterations in metabolic, epigenetic, and endocrine signaling pathways. Importantly, these hepatic findings parallel the divergent transcriptomic responses observed in the placenta, underscoring the placenta–liver axis as a central mediator of growth and metabolic programming following preconception phthalate exposure.

The contrasting effects of phthalate exposure on fetal versus adult growth may reflect a classic example of developmental reprogramming described by the Developmental Origins of Health and Disease (DOHaD) framework. At mid-gestation, increased fetal weight likely results from altered placental nutrient transfer and compensatory endocrine activity, consistent with the expansion of the junctional zone in male placentas and the upregulation of metabolic pathways in female placentas. These shifts may transiently enhance nutrient delivery and promote rapid growth in utero. However, such adaptations can create long-term tradeoffs. Dysregulated hepatic pathways in offspring, including excessive bile acid synthesis (*Cyp7a1*), altered fatty acid metabolism (*Cyp4a10*), disrupted one-carbon metabolism (*Mthfr*), and impaired immune/endocrine signaling (*Adm2, Ly6a*), are indicative of reduced metabolic flexibility postnatally. Over time, this mismatch between early placental-driven overnutrition and later hepatic dysfunction likely contributes to impaired nutrient utilization, antagonism of growth hormone/IGF signaling, and ultimately the reduced adult bodyweights observed in both sexes. This trajectory parallels the “Barker hypothesis,” which posits that perturbations in the intrauterine environment can lead to a thrifty phenotype: short-term survival advantages during development that predispose offspring to long-term growth restriction and metabolic disease.

Importantly, these mechanistic insights align with human epidemiological evidence. Several studies have reported associations between maternal phthalate exposure and altered birthweight, with outcomes ranging from fetal overgrowth to growth restriction depending on timing, dose, and mixture composition. For example, prenatal phthalate exposure has been linked to changes in placental size and function^96,97^ as well as increased risk of preterm birth^128^. Longitudinal cohort studies have further demonstrated that maternal and paternal preconception exposures predict reduced birth size and altered childhood growth trajectories.^84,129^ These findings parallel the growth patterns observed in our murine model, suggesting that the placenta–liver axis may represent a conserved mechanism of endocrine-metabolic reprogramming across species. Collectively, this evidence highlights the translational relevance of our work and underscores the need to consider preconception exposures in assessing the intergenerational impacts of environmental chemicals on human health.

### Conclusions and Limitations

Collectively, these findings demonstrate that preconception exposure to a human-relevant phthalate mixture disrupts maternal endocrine function, alters placental structure and transcriptome, and impairs offspring growth in a sex-specific manner. Female placentas exhibited broad transcriptional reprogramming across immune, metabolic, and structural pathways, whereas male placentas displayed structural changes linked to downstream developmental deficits. These sexually dimorphic responses underscore the importance of including both sexes in developmental toxicology research.

Our results reinforce the placenta’s role as a sensitive and integrative target of environmental exposures, mediating both immediate and long-term effects on offspring health. Moreover, they highlight the preconception window as a critical, yet often overlooked, period of vulnerability in the context of the developmental origins of health and disease (DOHaD).

Several limitations should be acknowledged. First, we utilized a single, environmentally relevant dose of a phthalate mixture, which limits our ability to assess dose-response relationships or define thresholds of effect. Second, we did not evaluate maternal or fetal tissue concentrations of parent compounds or metabolites, which restricts interpretation of internal exposure and pharmacokinetic variability. Future studies incorporating internal dosimetry and multiple dose levels would strengthen mechanistic inferences and relevance to human risk assessment. Finally, while CD-1 mice provide a robust outbred model, species-specific differences in placentation and hormone regulation may limit extrapolation to humans.

Despite these limitations, our study provides novel insights into how preconception exposures can shape reproductive outcomes and placental biology in a sex-specific manner. These findings emphasize the need to expand regulatory and research frameworks to include preconception as a critical window of exposure and support continued investigation into the long-term health implications of low-dose chemical mixtures.

## Acknowledgments

This project was funded by Clemson University Department of Biological Sciences Start-Up Funds (KDR). The authors thank Dr. Jodi Flaws and Mary Laws for providing guidance and protocols for preparation of the phthalate mixture. We would like to thank the Godley-Snell Research Center, particularly Tina Parker and Alison Walker, for their assistance with animal care and husbandry. Finally, we would like to thank the Clemson Light Imaging Facility, specifically Avery Herren for his assistance in processing and imaging placentas for histological assessment.

## Author contributions

Conceptualization: M.A. and K.D.R. Methodology: M.A. and K.D.R. Investigation: M.A., A.C.E., P.C.P., M.E.M, M.C.B., A.P.S., M.A.W., Z.J.P., and K.D.R. Visualization: M.A., A.C.E., and K.D.R. Funding acquisition: K.D.R. Project administration: K.D.R. Supervision: K.D.R. Writing-original draft: M.A., and K.D.R. Editing and Reviewing: M.A., A.C.E., P.C.P., M.E.M, M.C.B., A.P.S., M.A.W., Z.J.P., and K.D.R

The authors have no conflicts of interest to disclose.

