## Supplemental Figure for "Exploratory Assessment of Preconception Phthalate Exposure on Fertility and Offspring Health in Mice"

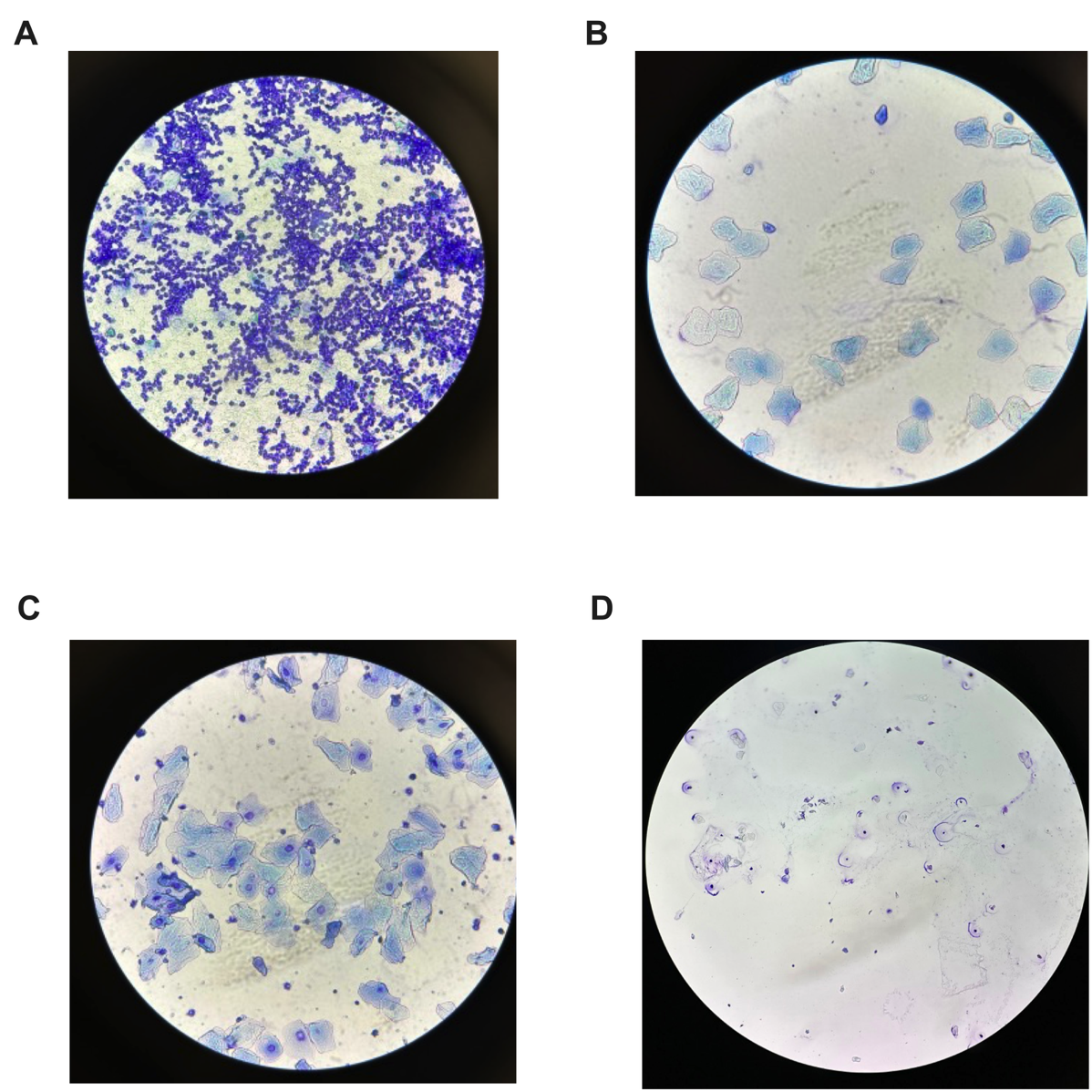

**Supplemental Figure 1**: ***Representative images of the stages of the Estrus cycle*. (**A) Proestrus stage: polymorphonuclear leukocytes B) Estrus stage: Cornified Epithelial cell C) Metestrus stage: Cornified Epithelial cells, polymorphonuclear leukocytes D) Diestrus stage: few numbers of mixed cell types, primarily leukocytes.

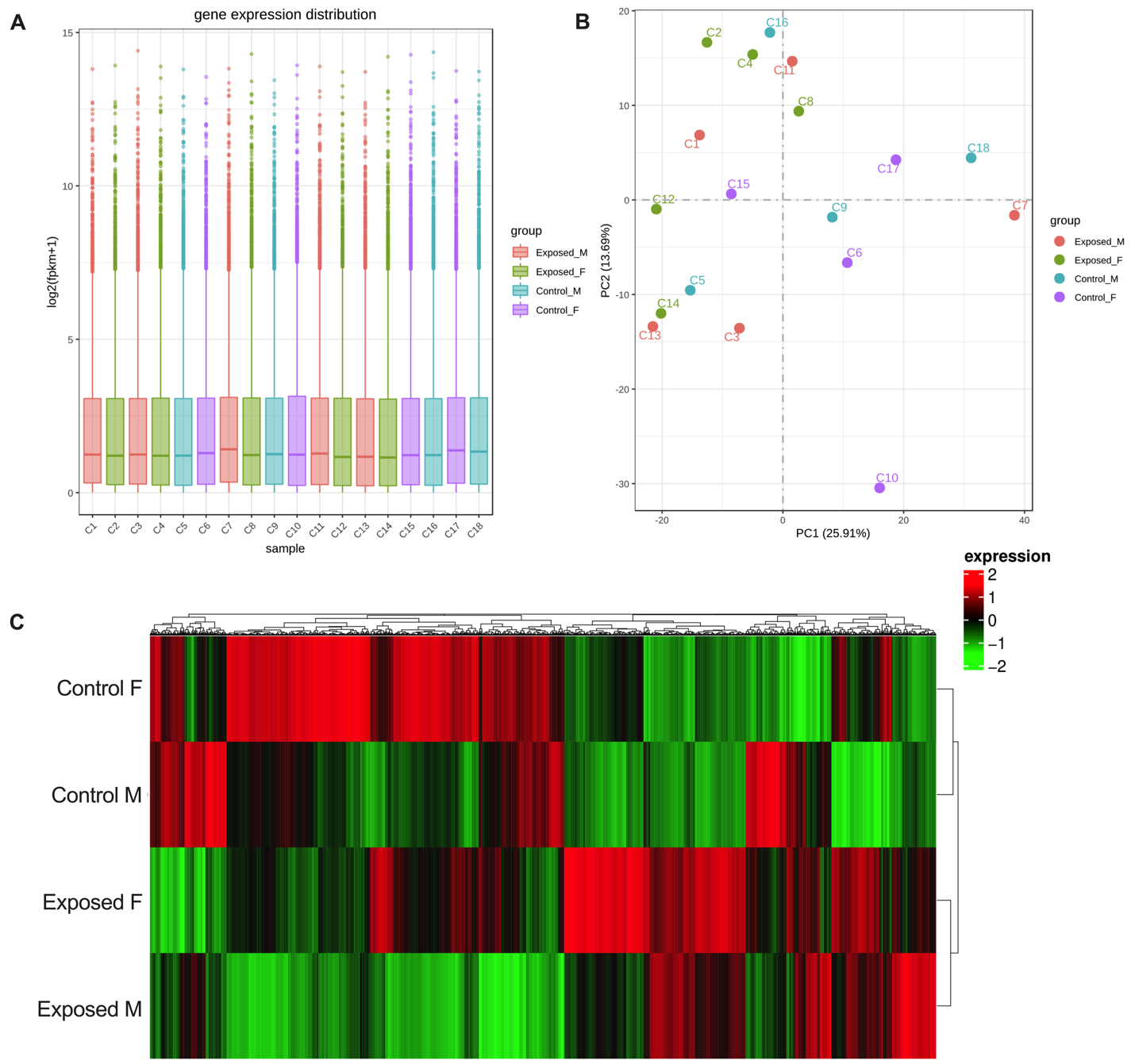

**Supplemental Figure 2**: (A) Gene expression distribution plot, (B) principal components analysis (PCA), and (C) heatmap of all the differentially expressed genes in male and female, control and exposed placentas.

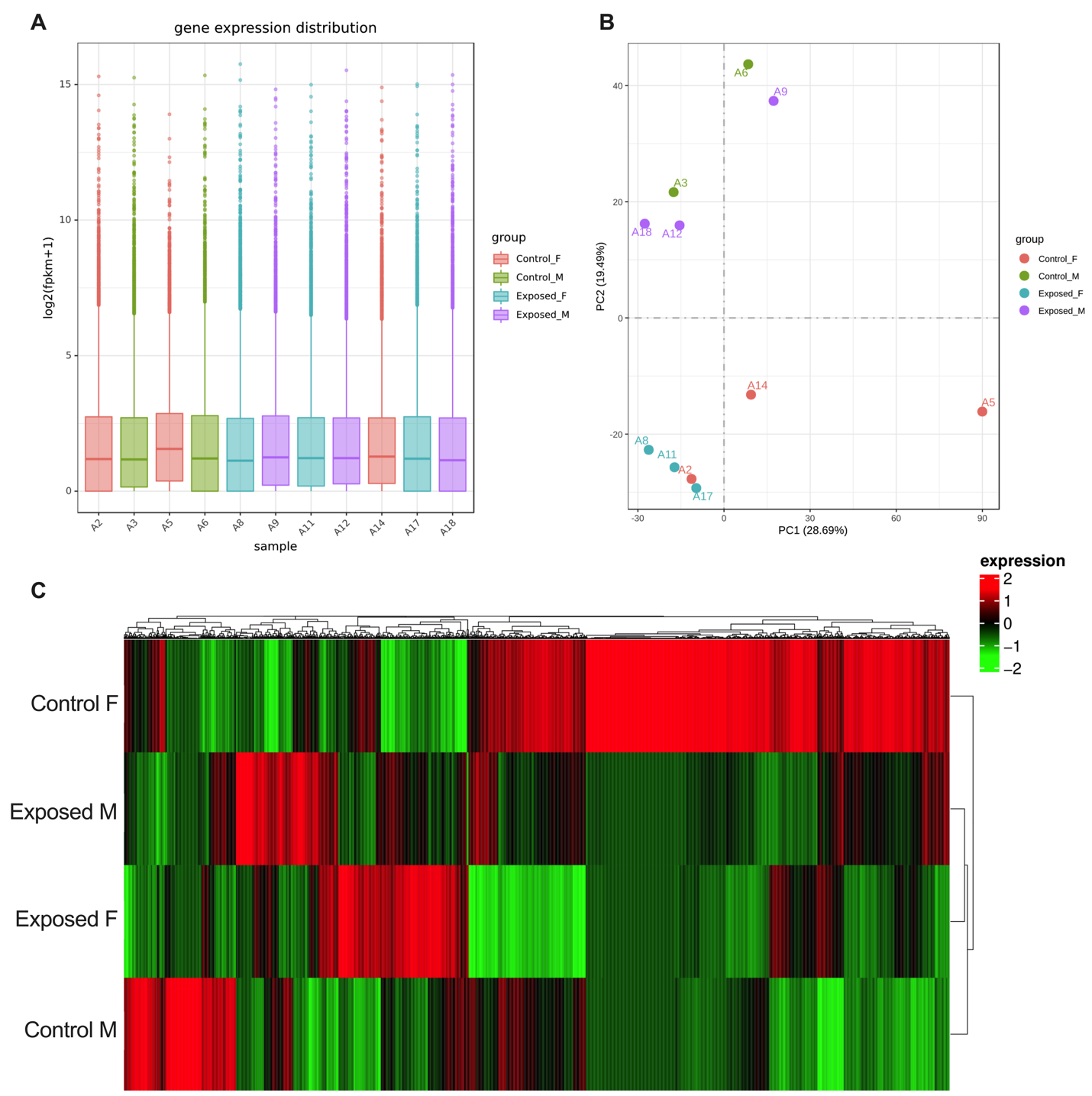

**Supplemental Figure 3**: (A) Gene expression distribution plot, (B) principal components analysis (PCA), and (C) heatmap of all the differentially expressed genes in male and female, control and exposed livers.
